# A mathematical investigation of chemotherapy-induced peripheral neuropathy

**DOI:** 10.1101/2020.04.23.057851

**Authors:** Parul Verma, Muriel Eaton, Achim Kienle, Dietrich Flockerzi, Yang Yang, Doraiswami Ramkrishna

## Abstract

Chemotherapy-induced peripheral neuropathy (CIPN) is a prevalent, painful side effect which arises due to a number of chemotherapy agents. CIPN can have a prolonged effect on quality of life. Chemotherapy treatment is often reduced or stopped altogether because of the severe pain. Currently, there are no FDA-approved treatments for CIPN partially due to its complex pathogenesis in multiple pathways involving a variety of channels, specifically, voltage-gated ion channels. A surrogate of neuropathic pain in an *in vitro* setting is hyperexcitability in dorsal root ganglia (DRG) peripheral sensory neurons. Our study employs bifurcation theory to investigate the role of voltage-gated ion channels in inducing hyperexcitability as a consequence of spontaneous firing, due to the common chemotherapy agent paclitaxel. Our mathematical investigation suggests that the sodium channel Na_v_1.8 and the delayed rectifier potassium channel conductances are the most critical for hyperexcitability in normal firing small DRG neurons. Introducing paclitaxel into the model, our bifurcation analysis predicts that hyperexcitability is extreme for a medium dose of paclitaxel, which is validated by multi-electrode array recordings. Our findings using multi-electrode array experiments reveal that the Na_v_1.8 blocker A-803467 and the delayed rectifier potassium enhancer L-alpha-phosphatidyl-D-myo-inositol 4,5-diphosphate, dioctanoyl (PIP_2_) have a protective effect on the firing rate of DRG when administered separately together with paclitaxel as suggested by our bifurcation analysis.

## Introduction

Chemotherapy-induced peripheral neuropathy (CIPN) is a painful, dose-limiting side effect of chemotherapy cancer treatment that affects more than 85% of patients during treatment [12] and 60% of patients three months post chemotherapy treatment [28]. These patients report enduring “pin and needle” paresthesias (i.e., tingling, numbness) in the peripheral nervous system (hands and feet) [24]. The onset of symptoms can range from one day to two years after treatment and can even persist throughout life, causing a significant decrease in the quality of life of cancer survivors [30, 5]. Furthermore, CIPN has been shown to result in depression, frustration, and a sense of loss of purpose among these patients [31]. To improve quality of life of cancer patients, it is imperative to find CIPN preventive agents. Since there is currently no FDA-approved treatment for CIPN, management of CIPN-induced pain includes many options such as antidepressants, anticonvulsants, anti-inflammatory, and opioid therapies.

Neurons in both the peripheral and central nervous system (PNS and CNS, respectively) are involved in the pain sensing and relay pathway. Although there is some role of the CNS in CIPN, treatments targeting CNS-based pain pathways have not been sufficient to reduce CIPN [7, 13]. Mainly, CIPN has been studied in the dorsal root ganglia (DRG) peripheral sensory neuron model. Since DRG are pseudo-unipolar, they can relay to other neurons in both the central and peripheral nervous system, thus allowing the different subpopulations of DRG to respond to different nociceptive stimuli including mechanical, thermal, and chemical [10]. Specifically, a nociceptive pain response can be identified through hyperexcitability, through increased spontaneous firing in voltage-gated sodium channels Na_v_1.6-1.9 [19] and potassium channels [9]. DRG are more susceptible to chemotherapy agents than the central nervous system neurons because DRG do not have an extensive neurovascular barrier to limit drug entry [18, 17]. Thus, we chose DRG to be our model system for this study. In particular, we focus on the mathematical model of a small DRG neuron, and validate results on a DRG neuron culture.

A myriad of alterations in various pathways have been linked to CIPN including voltage-gated ion channels, in calcium signaling, in fast axonal transport, and in occurrence of oxidative stress and inflammation have all been interlinked with CIPN [4]. Several chemotherapy agents, such as vincristine, paclitaxel, and oxaliplatin have been suggested to cause CIPN [4]. In this study, we focus on the chemotherapy agent paclitaxel (brand name Taxol). Paclitaxel is a microtubule-binding cancer agent used for several solid tumor cancers such as breast, ovarian, and lung. The conjecture concerning the paclitaxel-induced CIPN (PIPN) mechanism is that it reduces axonal transport of mRNA which can lead to axonal degeneration, alters expression of membrane ion channels, and induces inflammation and oxidative stress [4]. Several agents have been tested in clinical trials, but their ability to prevent PIPN is still unclear [1]. Since it can be difficult to control and examine multiple events in an experimental setting, a mathematical model can be employed to investigate mechanisms by exploring many channels in a controlled manner to observe how the system behavior changes upon any external influences. In this study, we analyze the role of voltage-gated sodium and potassium ion channels in paclitaxel-induced hyperexcitability using mathematical modeling and bifurcation theory, then support our results using *in vitro* recordings.

## Results

### Model description and simulations

We analyze a mathematical model representative of a small DRG neuron, which is a pain-sensing neuron. The model consists of the two sodium channels (Na_v_1.7 and Na_v_1.8), two potassium channels (delayed rectifier (KDR) and A-type transient (KA)), and one leak channel. We included these channels as they were deemed to be most prominent in small DRG neurons [14, 29, 6] and we wanted to investigate a minimal model to begin with and present a framework that can be suitably extended to more detailed models. The main equation for membrane voltage is written as:

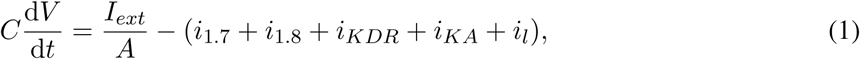

where, *I*_*ext*_ is the external applied current, *i*_1.7_, *i*_1.8_, *i*_*KDR*_, *i*_*KA*_, *i*_*l*_ are specific ionic currents due to Na_v_1.7, Na_v_1.8, delayed rectified potassium, A-type transient potassium, and leak channels. *C* is the specific membrane capacitance, *A* is the area, *V* the membrane voltage, and *t* is time. These ionic currents are written as following:

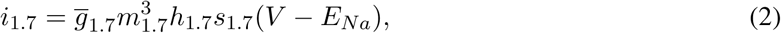

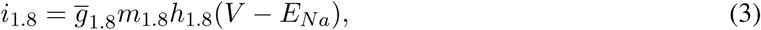

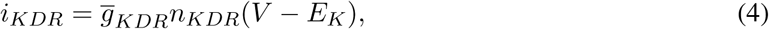

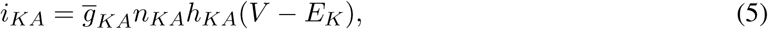

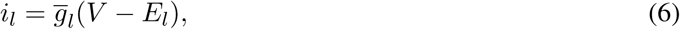

where, 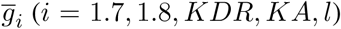 are maximal conductances and are constants. *E*_*Na*_, *E*_*K*_, and *E*_*l*_ are equilibrium ion potentials. All the activation and inactivation gating variables *x* (*x* = *m*_1.7_, *h*_1.7_, *s*_1.7_, *m*_1.8_, *h*_1.8_, *n*_*KDR*_, *n*_*KA*_, *h*_*KA*_) are written as the following in the Hodgkin-Huxley form [15]:

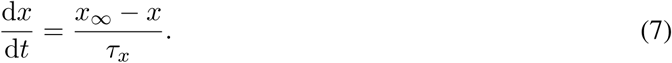

The expressions of *x*_∞_ and *τ*_*x*_, and the parameter values are specified in the Materials and methods section. All the equation forms have been extracted from literature [6].

An indicator of peripheral neuropathy is spontaneous firing: repetitive firing of action potentials for *I*_*ext*_ = 0. In this work, we explore the parameters that can lead to spontaneous firing and can be potentially impacted by paclitaxel. We vary the maximal ion conductances to explore whether they can induce spontaneous firing since current literature indicates that paclitaxel can manipulate the expression of voltage-gated ion channels [38]. A bifurcation analysis of this model with *I*_*ext*_ as the bifurcation parameter can be found elsewhere [32].

First, we performed dynamic simulations for different parameter values. The initial conditions correspond to the stable steady state solution obtained for the parameter values mentioned in Table 2. Fig 1 demonstrates how the voltage dynamics vary upon increasing 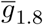 (We chose 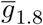for illustration purposes). For a low value of 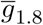, only a single action potential is observed and the system settles down to a steady state, shown in Fig 1A. For a higher value of 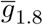, mixed-mode oscillations (MMOs) are observed, shown in Fig 1B. MMOs consist of both small amplitude (subthreshold) and large amplitude (action potential) oscillations. For a higher value, continuous firing of action potentials is observed, shown in Fig 1C. In the next section, we focus on the switch from steady state to continuous firing by treating different channel conductances as bifurcation parameters. Mixed-mode oscillations were investigated in detail in another work [33] (submitted).

**Table 1:** Spontaneous neurons. Pax: Paclitaxel, Na_v_1.8-: Na_v_1.8 blocker, KDR+: KDR enhancer

| Treatment | Paclitaxel (nM) | Fold change | Percent change |
| --- | --- | --- | --- |
| Media | None | 0.94 | -3.64 % |
| Pax | 250 | 1.42 | 21.09% |
| Na<sub>v</sub>1.8- | 250 | 1.27 | 11.83% |
| KDR+ | 250 | 1.28 | 5.64% |

**Table 2:** Model parameter values

| Parameter | Value | Units | Reference |
| --- | --- | --- | --- |
| $A$ (area) | 2168.00 | $\mu\text{m}^2$ | [6] |
| $C$ | 0.93 | $\mu\text{F}/\text{cm}^2$ | [6] |
| $E_{Na}$ | 67.10 | mV | [6] |
| $E_K$ | -84.70 | mV | [6] |
| $E_l$ | -58.91 | mV | [6] |
| $\bar{g}_{1.7}$ | 18.00 | $\text{mS}/\text{cm}^2$ | [6] |
| $\bar{g}_{1.8}$ | 7.00 | $\text{mS}/\text{cm}^2$ | [32] |
| $\bar{g}_{KDR}$ | 4.78 | $\text{mS}/\text{cm}^2$ | [6, 32] |
| $\bar{g}_{KA}$ | 8.33 | $\text{mS}/\text{cm}^2$ | [6, 32] |
| $\bar{g}_l$ | 0.0575 | $\text{mS}/\text{cm}^2$ | [6] |
| IC50 | 500 | nM | [26, 25] |
| $h_n$ | 1 | Unitless | Assumed |
| $\bar{G}_{Na,max}$ | 100 | $\text{mS}/\text{cm}^2$ | Assumed |
| $\bar{G}_{K,min}$ | 0.1 | $\text{mS}/\text{cm}^2$ | Assumed |

**Figure 1:**
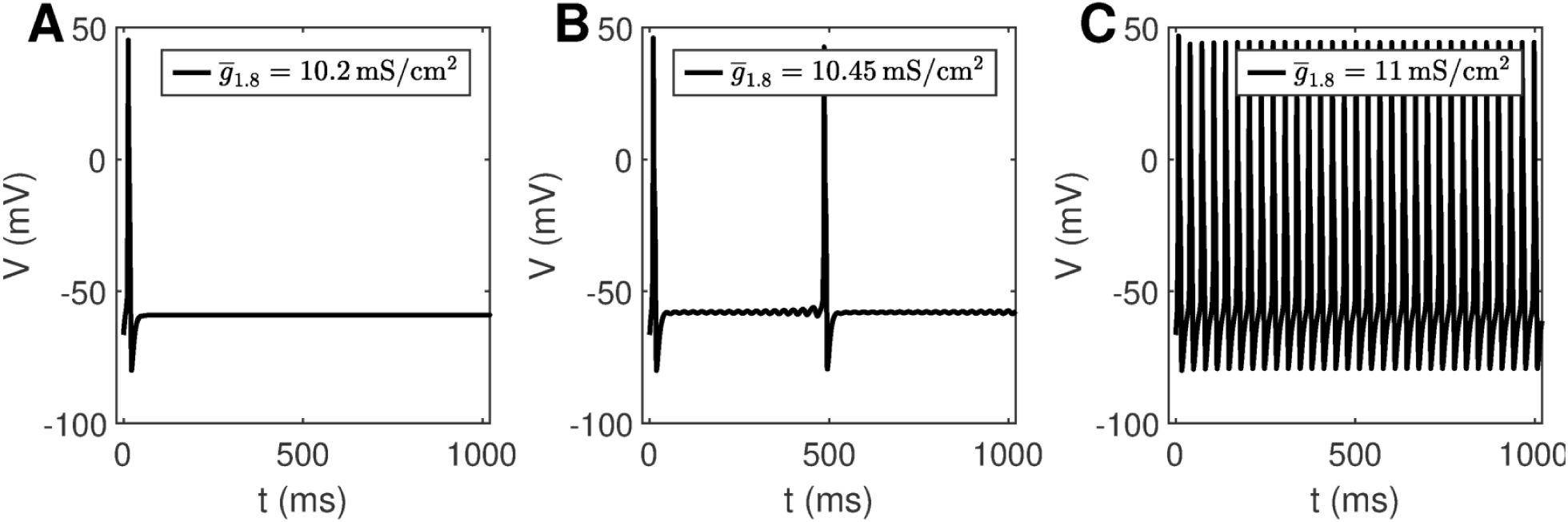
Dynamic simulations obtained by varying 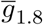. A: One action potential followed by a steady state is observed for 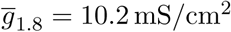, B: MMOs are observed for 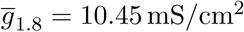, and C: Continuous firing of action potentials is observed for 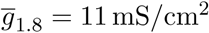

### Bifurcation analysis

XPPAUT is used to perform preliminary bifurcation analysis, and confirmation of results is done using MATCONT. Parameter settings for XPPAUT and MATCONT are mentioned in the Materials and methods section. As mentioned before, bifurcation diagrams are generated by setting the maximal conductance as the bifurcation parameter, one by one, for each voltage-gated ion channel involved in this model (Na_v_1.7, Na_v_1.7, KDR, and KA).

### One-parameter continuation

Firstly, we perform one-parameter continuations to find bifurcation points which could separate steady state from MMOs, and MMOs from continuous firing of action potentials. A partial bifurcation diagram is shown in Fig 2. As seen in Fig 2A and D, no bifurcation points are generated upon varying 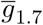 and 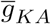. A single red line, representing stable steady state solutions, is observed. On the contrary, bifurcation points are observed in Fig 2B and C. In Fig 2B, a subcritical Hopf bifurcation point (HB) is detected upon increasing 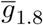. Beyond this point, steady state solutions become unstable, shown by the black branch. Two turning points/limit points (LP_1_, LP_2_) are also detected on the black branch of unstable steady state solutions. Since the Hopf bifurcation point is subcritical, unstable periodic solutions emanate from it, as shown by the blue branch. This branch first turns at a cyclic limit point (CLP_1_) and then meets the unstable steady state branch, indicating a homoclinic orbit. This turning is not obvious from the figure, however, it can be observed upon zooming into the branch. Upon moving in the backward direction starting from a large value of 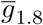 resulting in stable periodic solutions, a stable periodic solution branch is generated, indicated in green, finally leading to a cyclic limit point (CLP_2_) beyond which the periodic solutions become unstable, indicated in blue. This unstable periodic branch abruptly ends due to the period of the branch increasing substantially, indicating that it may tend towards a period-infinity solution, as shown in Fig 2E. The stable periodic solution branch indicates the spontaneous firing parameter regime. The frequency of spontaneous firing increase with 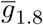, as shown in Fig 2E. MMOs are found in some region between the Hopf bifurcation point (HB) and the cyclic limit point (CLP_2_), shown by the shaded pink region. A detailed discussion for a similar situation is given in [33] (submitted).

**Figure 2:**
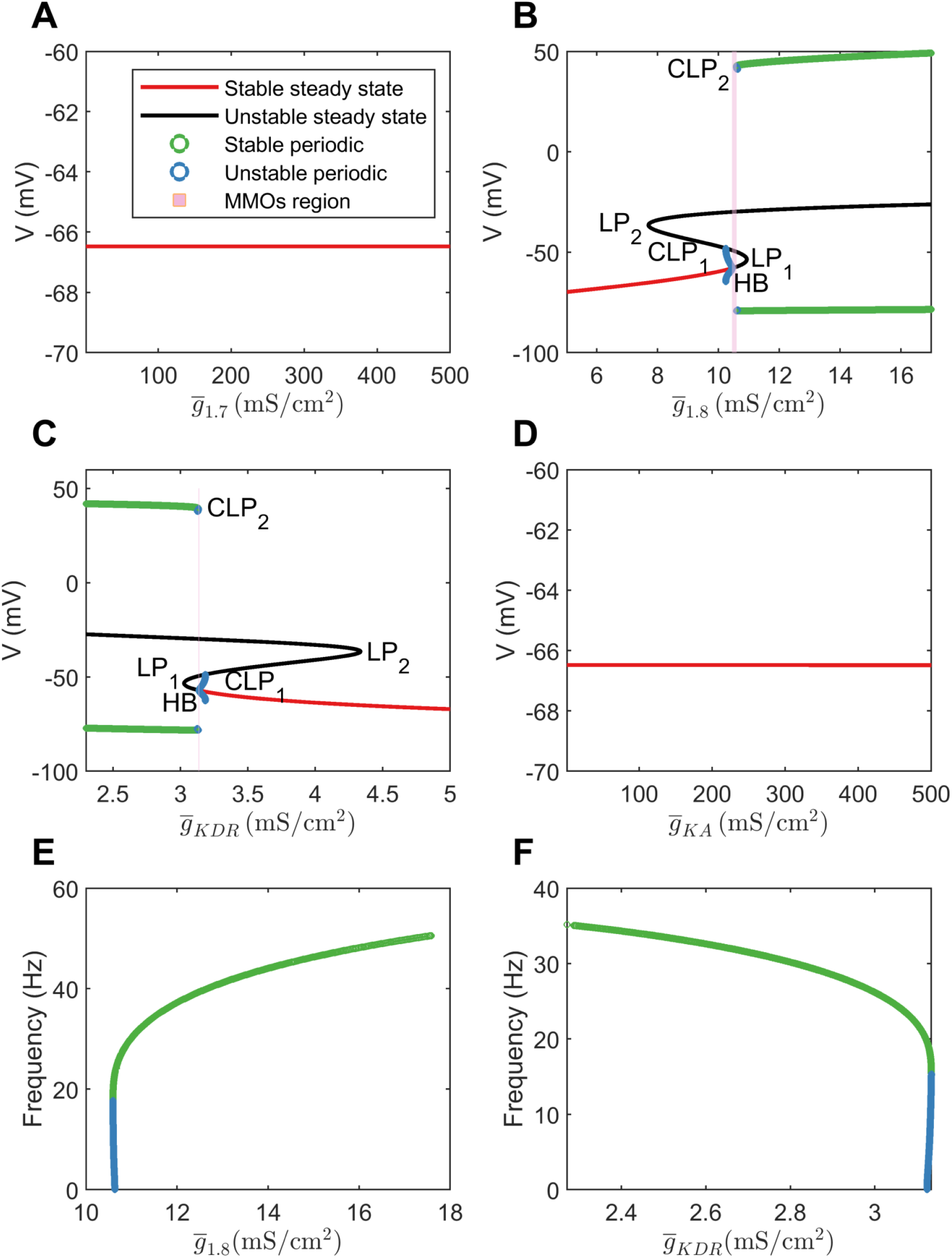
A-D: Bifurcation diagrams obtained by keeping 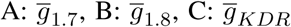, and 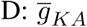 as the bifurcation parameters. E-F: Frequency versus maximal conductance obtained in the periodic firing regime with 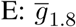 and 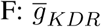 as the bifurcation parameters. The frequency of firing increases with 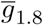 and decreases with 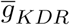. The frequency of unstable periodic solutions tends towards zero, implying that the unstable branch is ending in a period-infinity solution.

A similar, although horizontally flipped, bifurcation diagram is generated with 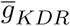 as the bifurcation parameter, shown in Fig 2C. Upon decreasing 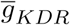, a subcritical Hopf bifurcation point (HB) is detected with unstable periodic solutions emanating from it. These unstable periodic solutions also intersect the unstable steady state branch, indicating a homoclinic orbit. Similarly to Fig 2B, a stable periodic solution branch (indicated by green circles) is detected as well which becomes unstable after a cyclic limit point (CLP_2_). As shown in Fig 2F, the unstable periodic solution again seems to tend towards a period-infinity solution. Moreover, the frequency of spontaneous firing decreases with increase in 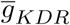.

These bifurcation diagrams indicate that manipulating 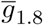or 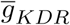 can induce spontaneous firing, while manipulating 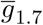 or 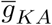 will not decrease hyperexcitability. Therefore, to reverse hyperexcitability, Na_v_1.8 and KDR channels should be targeted. We targeted these channels in case of paclitaxel-induced hyperexcitability, in a DRG culture, the results of which are described in the Experimental validation results section.

### Two-parameter continuation

We also perform two-parameter continuation in order to explore the combinational effects of these conductances. To this end, we observe the changes in the detected bifurcation points upon changing another maximal conductance. In particular, we perform continuation of HB, LP_1_, LP_2_, and CLP_2_. These results are shown in Fig 3.

**Figure 3:**
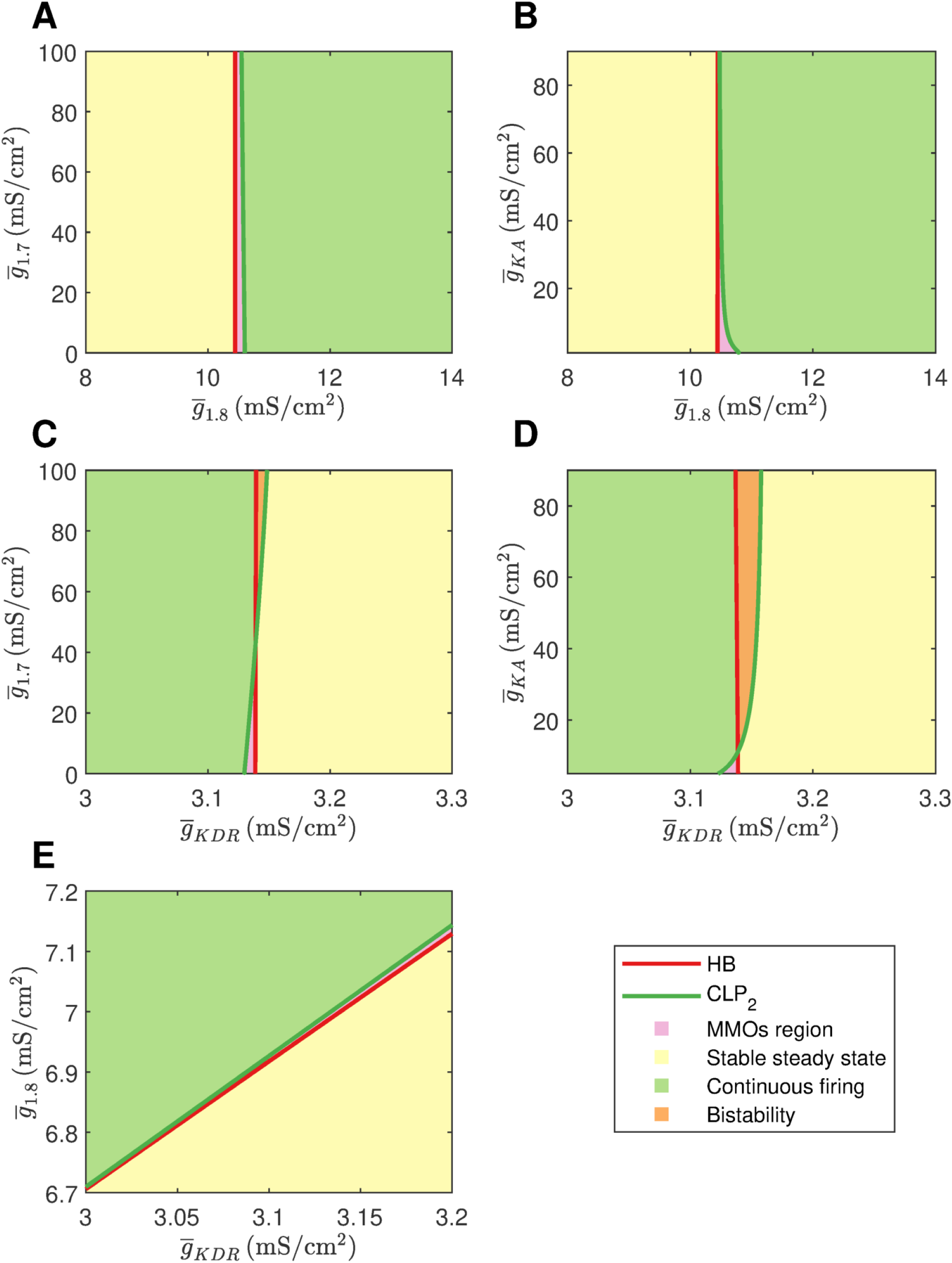
Two parameter continuations performed for the Hopf bifurcation point HB, limit points LP_1_ and LP_2_, and cyclic limit point CLP_2_. A: Continuation plot for 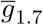 versus 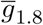 show that the bifurcation points generated by keeping 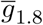 as the bifurcation parameter do not shift upon varying 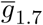. B: HB and LP’s of 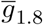 bifurcation diagram do not shift upon varying 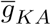. CLP_2_ shift rightwards upon decreasing 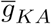. This implies that the MMOs region will be wider in this case. C: Bifurcation points of 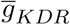 do not shift upon varying 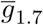. D: HP and LP’s of 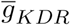 do not shift upon varying 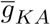. CLP_2_ shifts leftwards upon decreasing 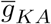. This implies that the MMOs region will become narrower in this case. E: A linear combinational effect is seen between 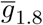 and 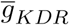. Note that the thin gap between stable steady state and continuous firing regimes is the MMOs region.

As seen in Fig 3A and 3B, 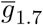 and 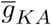 do not impact the bifurcation points substantially even in combination with 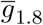. Decrease in 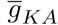 can shift the cyclic limit point of 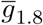 to the right, as seen in Fig 3B, which implies that the MMOs regime will become wider. In both the cases, the bifurcation points vary within a narrow range of 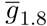. Similarly, Fig 3C and 3D show that 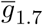 and 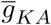 do not impact the bifurcation points substantially even in combination with 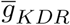. In these two cases, increase in 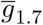and 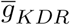 can lead to a region of bistability where stable steady state and continuous firing of action potentials solutions coexist. However, the bifurcation points only vary within a narrow range of 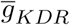. Upon varying 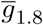 and 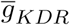 together, the bifurcation points vary linearly, as shown in Fig 3E. This indicates that decreasing 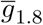 and increasing 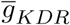 can eliminate spontaneous firing.

### Effect of paclitaxel

Current literature suggests that paclitaxel can impact gene expression of various voltage-gated ion channels [38, 2]. However, it is not known whether the impact is direct or indirect. Evidence indicates that paclitaxel impacts inflammatory cytokines, which can subsequently manipulate the ion channels [2]. For example, these inflammatory signals can increase sodium current [34]. Moreover, a sigmoidal dose-dependent relation is observed between paclitaxel and macrophage IL-12, as seen in Fig 3 in [22]. Based on these evidences, we assumed a Hill’s kinetics type relation between paclitaxel and ion channel maximal conductances. Hill’s kinetics are widely used to model dose-response curves. We assume that the conductances will vary as a function of paclitaxel dosage. Moreover, we assume that paclitaxel will lead to an increase in maximal conductance of both the sodium channels, while it would lead to a decrease in maximal conductance of both the potassium channels since these cases would lead to spontaneous firing. We assume that all the conductances get impacted even though all of them may not lead to a change in spontaneous firing, as shown in the previous section. Therefore, we consider the following relationships:

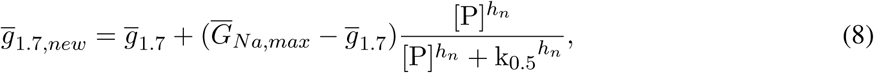

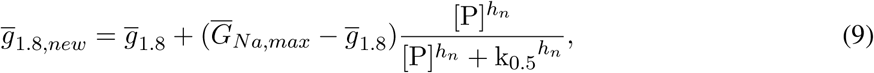

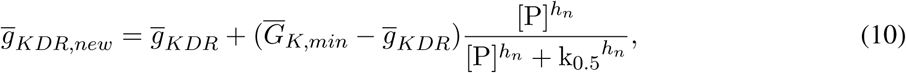

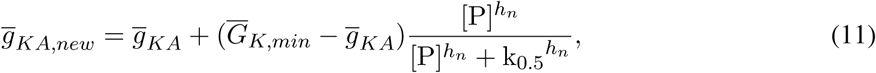

where, *h*_*n*_ is Hill’s coefficient, [P] is the paclitaxel concentration (in nM), k_0.5_ is the half maximal effective concentration, 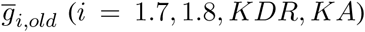 is the original maximal conductance value, 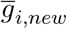 is the updated maximal conductance value from the above equation. 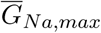 and 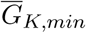 stand for the upper and the lower limit of the maximal conductances, respectively.

Depending on 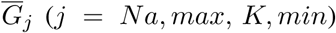 and *h*_*n*_, the relationship between [P] and 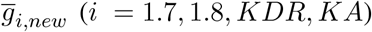 will vary, as shown in Supplementary figure S1 FigA, S1 FigB, and S1 FigC. Increasing or decreasing 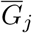 increases or decreases the maximal conductance parameter 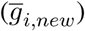 range covered upon varying [P]. Large *h*_*n*_ creates sigmoidal curves. Decreasing *h*_*n*_ makes the curve seem exponential. The values of 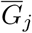 and *h*_*n*_ are assumed. See supplementary figure S2 Fig for a sensitivity analysis of these parameters with respect to the solution regimes.

Following this, we perform numerical bifurcation analysis with paclitaxel concentration [P] as the bifurcation parameter. A partial bifurcation diagram is shown in Fig 4A. Fig 4A shows that upon stable steady state solution continuation, a subcritical Hopf bifurcation point (HB_1_) is found beyond which the solutions become unstable. Unstable periodic solutions emanate from this Hopf bifurcation point. Upon continuation of the unstable steady state solution branch in black, a supercritical Hopf bifurcation point (HB_2_) is found, beyond which the steady state solutions become stable again. Stable periodic solutions (indicated in green) emanate from this point. The paclitaxel-interval between these two Hopf bifurcation points constitutes the spontaneous firing regime. The subinterval where the periodic and the steady state branch are unstable (between PD and CLP_4_) corresponds to the MMOs regime. MMOs of this model have been studied in some detail in [33] (submitted); a detailed investigation of the MMOs shown in Fig 4A was beyond the scope of this work. Stable periodic solutions with a small amplitude arise from the supercritical Hopf bifurcation point which turn unstable after a cyclic limit point (CLP_4_). The unstable periodic solutions finally become stable after a subcritical period doubling bifurcation point (PD) when going in the direction of decreasing [P]. The stable periodic solutions become unstable again after a cyclic limit point (CLP_2_).

**Figure 4:**
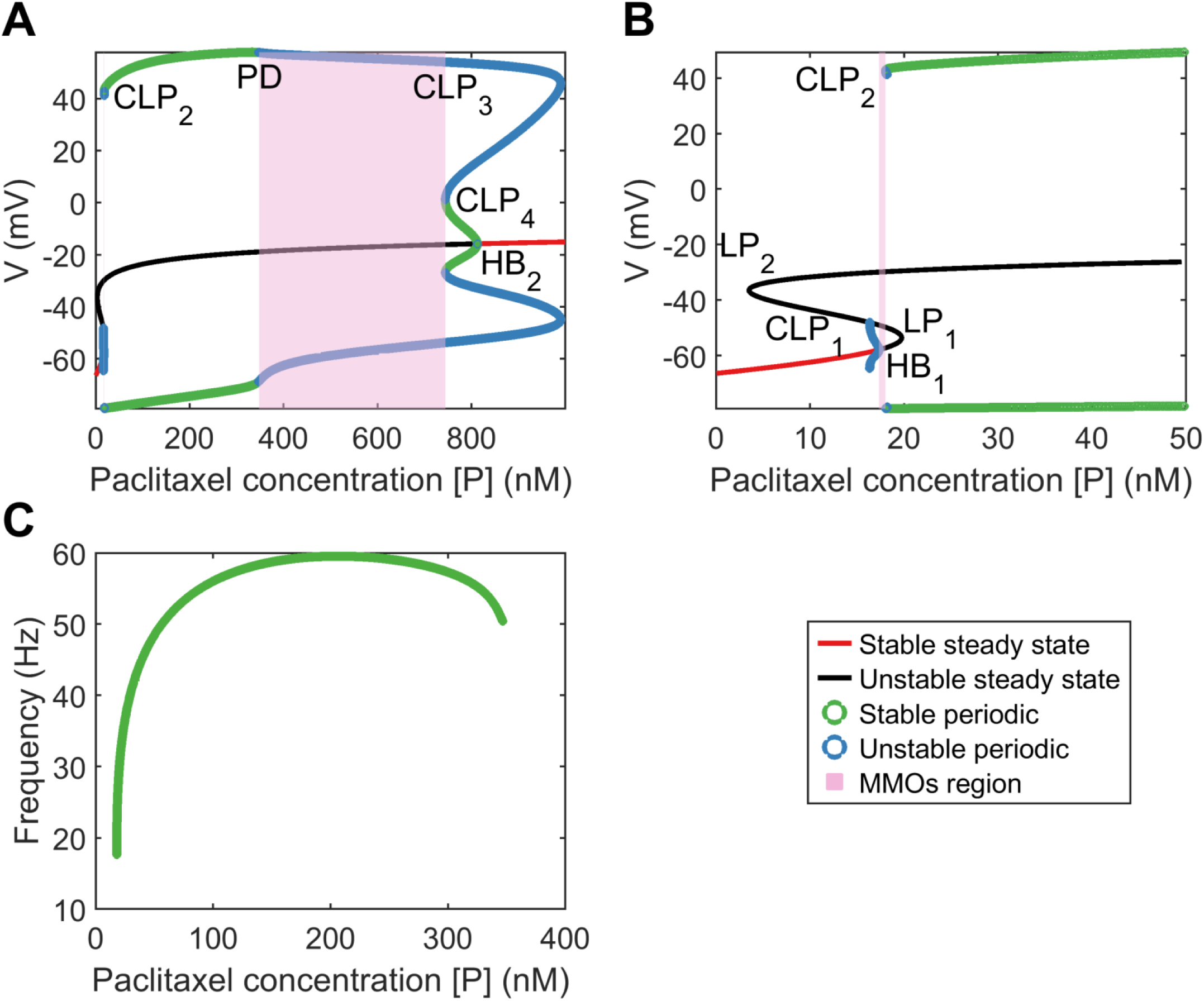
A: Bifurcation diagram obtained by treating paclitaxel concentration as the bifurcation parameter. B: A zoomed in version of the bifurcation diagram in A. HB_1_: subcritical Hopf bifurcation point, HB_2_: supercritical Hopf bifurcation point, LP_1_ and LP_2_: limit points, CLP_1_, CLP_2_, CLP_3_, and CLP_4_: cyclic limit points, PD: periodic doubling bifurcation point. C: Frequency plot for the stable periodic firing region. Frequency first increases and then decreases upon increasing paclitaxel concentration. Left and right end points of this curve refer to CLP_2_ and PD, respectively.

The frequency of firing in the first stable periodic solutions regime (between CLP_2_ and PD) is shown in Fig 4B. It is shown that upon increasing paclitaxel concentration, frequency of firing first increases, and then decreases after reaching a maximum firing rate. Beyond the PD point, the frequency of firing decreases further since the solutions are of MMOs type.

### Experimental validation results

A high-throughput way to continuously record neuronal firing patterns is to use multielectrode array (also referred to as microelectrode array, MEA). This system is capable of recording of extracellular voltage potentials with millisecond temporal resolution of neurons in cultures grown on a 768 array of electrodes up to 96 well format, which makes it high-throughput.

To validate the mathematical modeling of the effect of paclitaxel dose on hyperexcitability, we measured the firing for different doses of paclitaxel. Low doses (10 nM) and high doses (1 *µ*M) of 24-hour paclitaxel administration caused lower firing than 250 nM paclitaxel as expected (Fig 5A). Thus, we decided to use 250 nM for the dose of the subsequent experiments. Na_v_1.8 blocker A-803467 and KDR enhancer L-alpha-phosphatidyl-D-myo-inositol 4,5-diphosphate, dioctanoyl (PIP_2_) when administered separately together with paclitaxel reduces the number of spontaneous firing neurons (Table 1) and firing rate (Fig 5B). Similarly, the representative heat maps of firing frequency reveals the same trend qualitatively (Fig 5C). Specifically, the mean firing rate fold change from baseline is 1.16 ± 0.07 for media control and increases to 1.50 ± 0.10 for paclitaxel (p<0.0001 compared to media). Na_v_1.8 blocker A-803467 decreases the firing rate to 1.28 ± 0.07 (p=0.0449 compared to paclitaxel). Similarly, KDR enhancer PIP_2_ reduced paclitaxel-induced hyperexcitability to 1.28 ± 0.09 (p<0.0001 compared to paclitaxel). These results match the prediction from the bifurcation analysis that blocking Na_v_1.8 and enhancing KDR will protect against paclitaxel-induced hyperexcitability.

**Figure 5:**
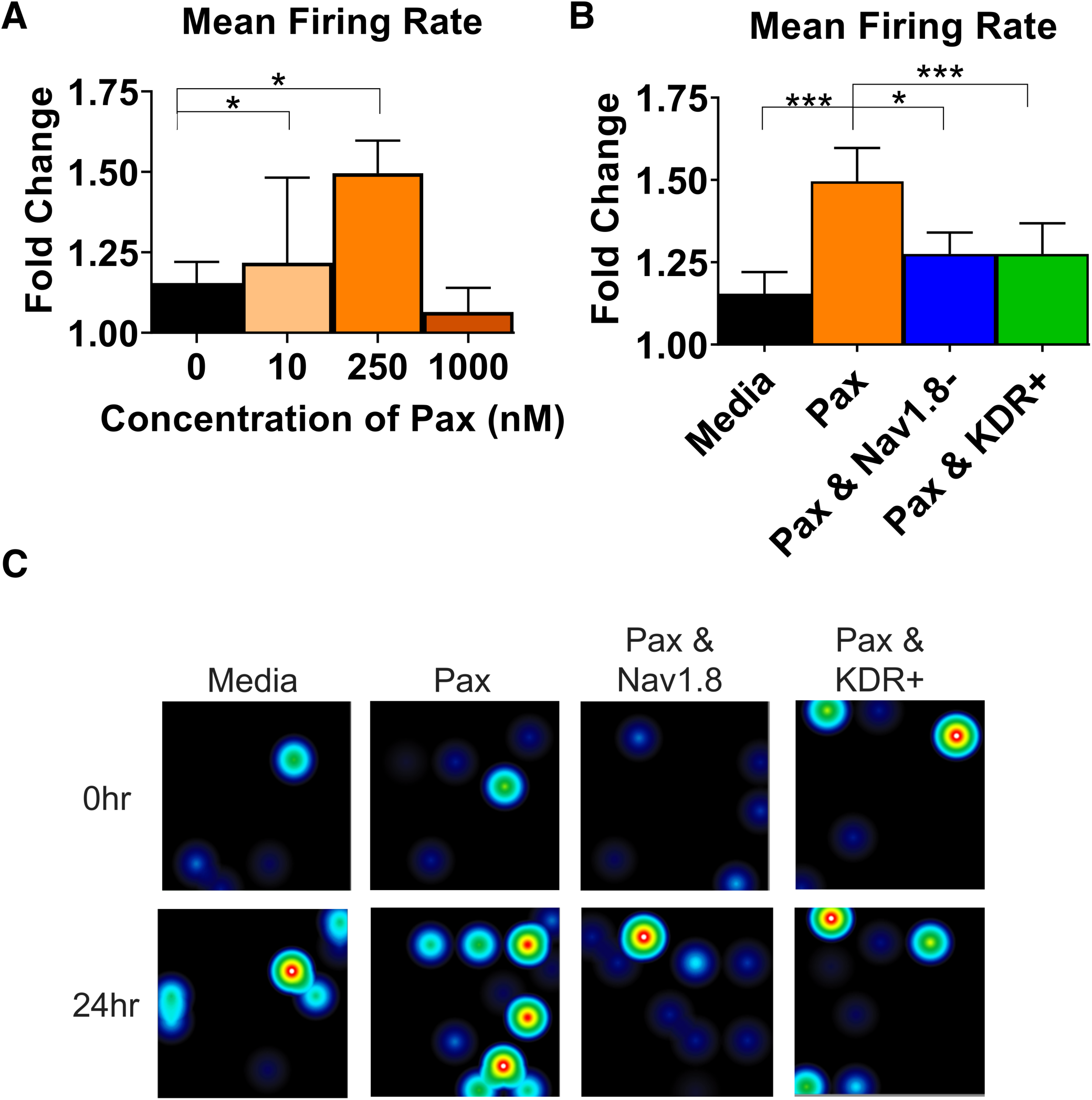
Multielectrode array (MEA) firing summary shows amelioration of hyperexcitability after treatment of A-803467 (Na_v_1.8 blocker) and PIP_2_ (KDR enhancer). All parameters are reported as fold change (treatment over baseline of culture before treatment). A) Mean firing rate for different dosages of paclitaxel. B) Mean firing rate reveals a significant increase in paclitaxel firing from media control (p<0.0001), a significant decrease from paclitaxel when A-803467 and PIP_2_ are administered separately (p=0.0449 and p<0.0001, respectively). C) Heatmap of representative MEA recordings with firing frequency of each active electrode color-coded: warm colors (red, orange, yellow) represent high firing frequency (white=10Hz); cool colors (green, blue) represent low firing frequency (black=0Hz). Each circle represents a spontaneous firing neuron within the 8 × 8 electrode array. Top row is baseline at time 0 before treatment is added. Bottom row is 24 hours after treatment was added. Asterisks denote statistical significance from Mann-Whitney U test (*P<0.05, **P<0.01, ***P<0.001).

**Figure 6:**
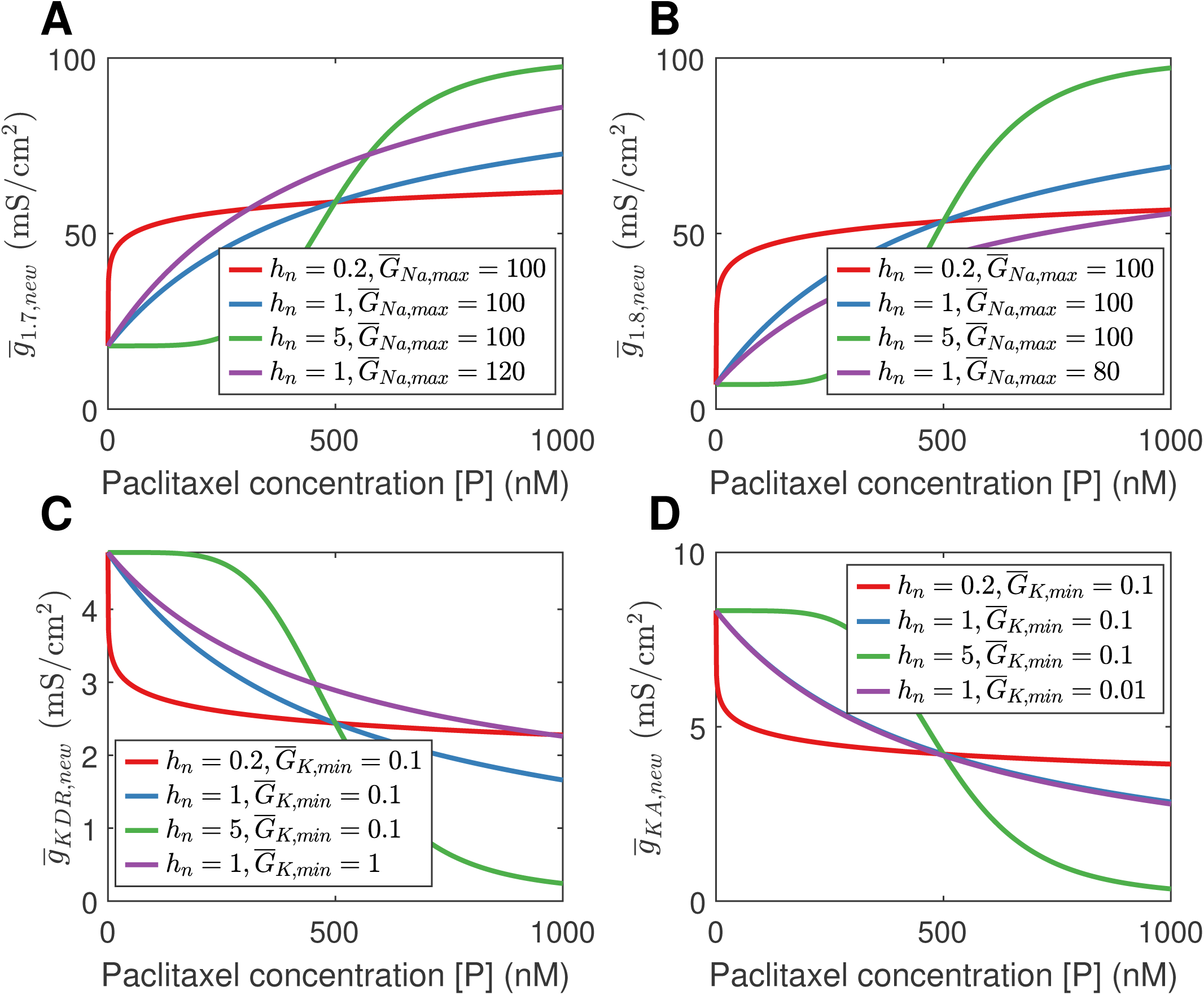
Effect of paclitaxel on conductances and firing. A: Effect of paclitaxel on 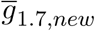, upon varying *h*_*n*_ and 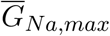. Increasing 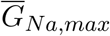 will widen the parameter range of 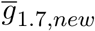,. Increasing *h*_*n*_ alters the curve to become more sigmoidal. B: Similar effect is seen with 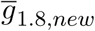. Decreasing 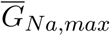 will reduce the parameter range of 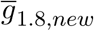. C: Increasing paclitaxel concentration decreases 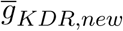. As before, increasing *h*_*n*_ makes the curve more sigmoidal. Decreasing 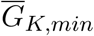 increases the parameter range. D: Similar effect is seen for 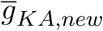. Increasing 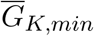 decreases the parameter range in this case. The blue curves correspond to the parameter values that were considered for bifurcation analysis. Note that the blue and purple curves are overlapping in in D, thus the blue curve is not visible.

**Figure 7:**
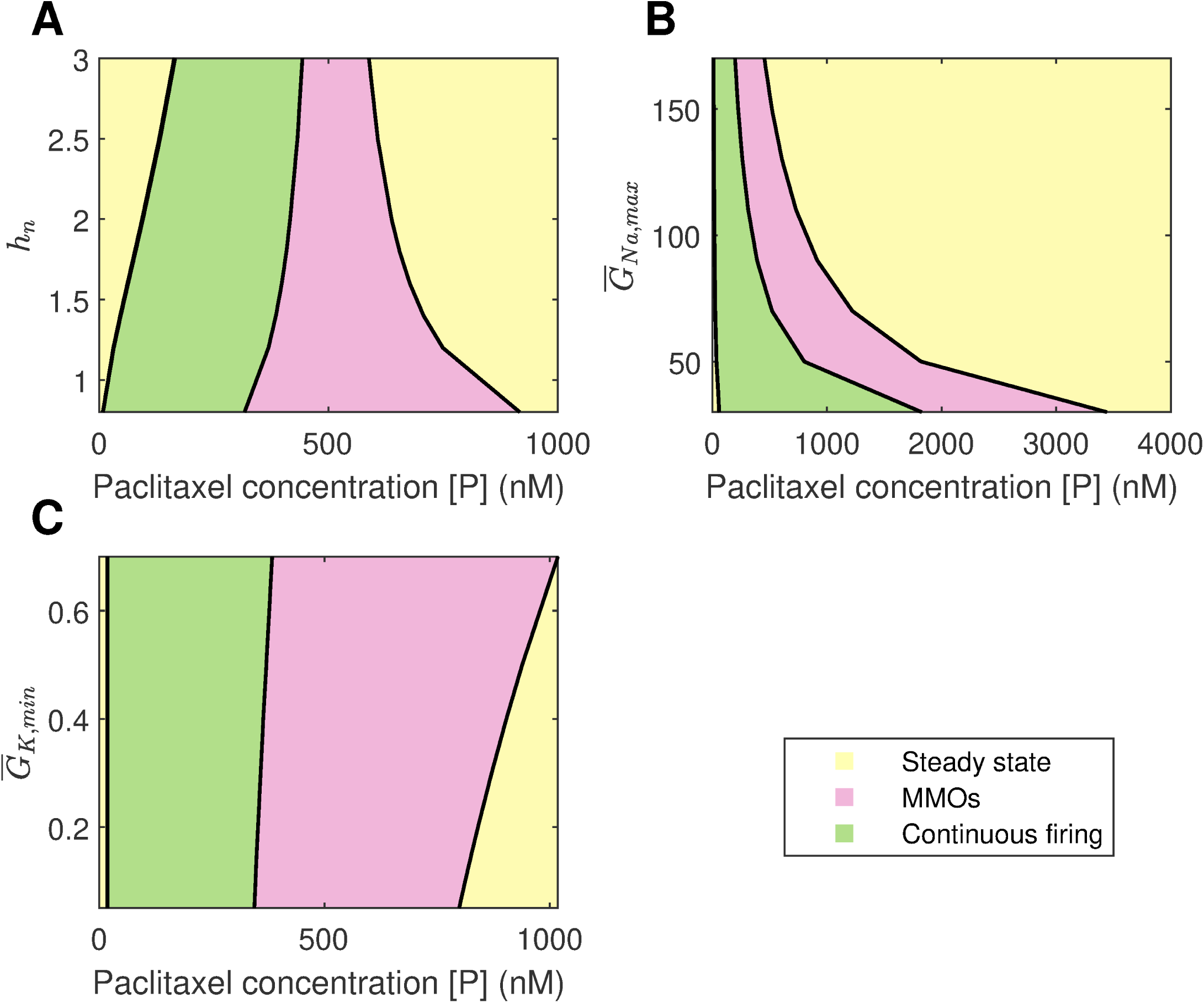
Regions of stable steady state, MMOs, and continuous firing upon varying *h*_*n*_, 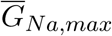, and 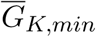 with paclitaxel concentration [P]. A: Continuation of Hill’s coefficient *h*_*n*_. Upon increasing *h*_*n*_, the spontaneous firing regime becomes narrower. B: Continuation of 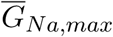. Upon increasing 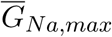 the spontaneous firing regime becomes narrower. C: Continuation of 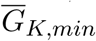. Upon increasing 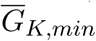, the spontaneous firing regime becomes wider.

## Discussions

CIPN is a debilitating experience for cancer patients with no current established methods of preventing or treating it due to minimal understanding of its pathophysiology [24]. In this work, we apply a novel mathematical approach using bifurcation theory to understand the role of sodium and potassium ion channels in CIPN. To this end, we analyzed a mathematical model representative of a small DRG neuron. Maximal conductances were kept as bifurcation parameters to identify those that can induce spontaneous firing (an indicator of peripheral neuropathy). We further used MEA experiments to support our findings.

Using bifurcation theory, we find that, increasing 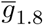 and decreasing 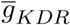 can induce spontaneous firing (see Fig 2). The effect may be aggravated in combination, as seen from the two parameter plot in Fig 3E. Our results indicate that a Na_v_1.8 blocker should reduce spontaneous firing which supports the role of Na_v_1.8 in contributing to the increased excitability in peripheral neuropathy [35, 39]. The significance of blocking Na_v_1.8 in PIPN was also observed in our MEA experiment in that Na_v_1.8 blocker A-803467 had a neuroprotective effect on paclitaxel-induced hyperexcitability when administered together in DRG neuron culture (Fig 5). Similarly, KDR was indicated in our model and MEA finding to be involved with hyperexcitability which is supported by literature [9]. Although these results are for specific dosages of A-803467 and PIP_2_ to observe the change in excitability, this electrophysiology data support the trends found in the bifurcation analysis. Depending on the amount of A-803467 and PIP_2_, hyperexcitability should change. Interestingly, Na_v_1.8 and KDR channels were found to be sensitive even with *I*_*ext*_ as the primary bifurcation parameter [33] (submitted). It will be of interest to investigate their protective effects on CIPN due to other chemotherapy agents such as vincristine and oxaliplatin, as well.

We assume a Hill’s type kinetics for the effect of paclitaxel on the ion channels. Based on this, a partial bifurcation diagram is generated, treating paclitaxel concentration as the bifurcation parameter. This bifurcation diagram indicates that spontaneous firing should arise for a mid-range of paclitaxel dosage. Moreover, firing rate should first increase and then decrease, seen from the frequency diagram in Fig 4C. A similar trend is seen in the MEA recordings, shown in Fig 5A. Firing rate is larger for the middle values of paclitaxel concentration in the range considered here. The frequency diagram (Fig 4C) may not reflect the actual reason for this trend. It may be that the cells died at a higher concentration, or they may not fire because of manipulation of the ion channels as shown by a mathematical relationship with paclitaxel. Further investigation is required to establish with certainty the reason behind this observation.

The parameters for the relationship between paclitaxel and ion channel maximal conductances are also assumed (see Table 2). In the supplementary figure S2 Fig, we have shown how the behavior of the model varies upon changing these parameters. Upon varying *h*_*n*_, 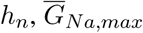, or 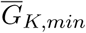, the qualitative behavior of the diagram does not change for the range of parameters considered here; stable steady states, MMOs, and continuous firing regimes are observed. Assuming that Hill’s kinetics is a reasonable relationship between paclitaxel and ion channel conductance, defined upper and lower limits of how much the conductances can vary may exist. These parameters can be estimated by recording currents due to each of these ion channels for different doses of paclitaxel, using patch-clamping experiments. Lastly, the value of k_0.5_ we used in our mathematical model (500 nM) is different from the paclitaxel dose amount used for the MEA experiment (250 nM). This is because we estimated the value from literature which was based on the IC50 value for iPSC-derived human neurons and clinical data based on paclitaxel’s toxicity on cancer cells [26, 25]. However, as seen in 4C, maximum spontaneous firing is observed around 200 nM. Since we only intend to perform a qualitative comparison between the bifurcation diagram (Fig 4A) and MEA paclitaxel dose trend (Fig 5A), the value of k_0.5_ assumed is not important. A different value of k_0.5_ will shift the complete bifurcation diagram, however, the qualitative structure of the diagram will remain the same.

The mathematical model that we considered is a minimal model representing dynamics of one type of peripheral neurons: small DRG neurons. *In vivo*, many more ion channels such as inward rectifier potassium and calcium channels are present in this neuron, and they need to be added to the model in future studies. More detailed mathematical models have been developed previously and can be used for this purpose [20]. Paclitaxel can also induce cytosolic calcium oscillations [3], which can again be analyzed using bifurcation theory. Another factor to add in future models is the impairment of axonal transport that occurs due to paclitaxel and its effect on microtubules [23], which can be modeled using cable equations [16]. As mentioned previously, the effect of paclitaxel on ion channels may be indirect and due to other inflammatory cytokines [2]. In the future, this can also be included in the model. Lastly, our model consists of only small DRG neurons. However, paclitaxel impacts medium and large DRG neurons as well [38]. It will be of interest to evaluate models representing different subtypes of DRG neurons and investigate the role of voltage-gated ion channels specific to each of them. The experimental setup recorded firing of all DRG subtypes. The blocker and enhancer may have regulated ion channels in other DRG neurons as well. However, we can be certain that Na_v_1.8 blocker acted on small DRG neurons since it is only present in them. PIP_2_ may have acted on other ion channels of other DRG subtypes. Similarly, paclitaxel may have impacted the firing rate of all DRG subtypes. This will require further investigation in the future by recording DRG subtype-specific firing.

DRG neuron firing has been correlated with neuropathic pain *in vitro* [36, 37], thus analyzing changes in firing patterns allows for a surrogate to study “pain in a dish”. Recent advances in neuroelectrophysiology technology have allowed for more temporal and spatial dynamic data collection on live cells. We chose a non-invasive method to measure the electrophysiological properties (spontaneous/non-evoked firing rate) of a network of neurons through microelectrode array (MEA) recording. Therefore, we used MEA to support our model outputs by recording the neuronal activity of the effect of paclitaxel on cultured DRG neurons, as shown in Fig 5.

Our bifurcation analysis identified that Na_v_1.8 and KDR are involved with the induction of spontaneous firing, associated with paclitaxel-induced hyperexcitability. This approach, along with patient-specific pharmacokinetics of paclitaxel, can be applied to the clinic by obtaining the patient’s electrophysiological profile using patch clamp or MEA and using those values as parameters in our model to determine the optimal dose of paclitaxel based on the individual, and for examining acute versus chronic onset of PIPN. Also, treatments can be designed to specifically block Na_v_1.8 or enhance KDR conductance to reduce hyperexcitability caused by paclitaxel since we have identified the corresponding channels as the most relevant ones.

## Materials and methods

### Model equations

Below, we describe the equations for each of the voltage-gated ion channels.

#### Na_v_1.7 equations

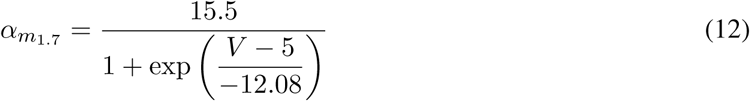

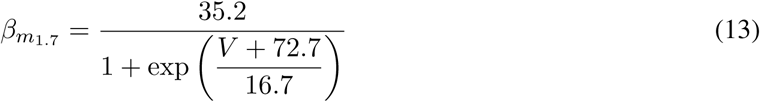

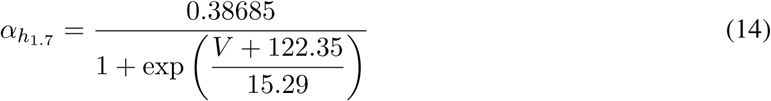

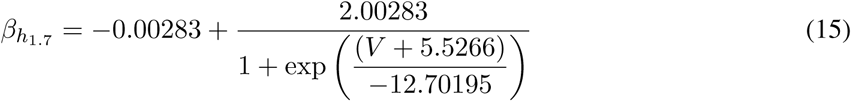

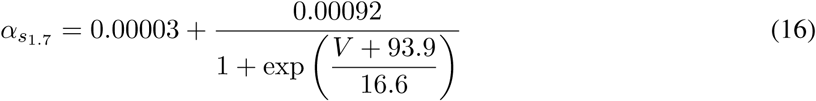

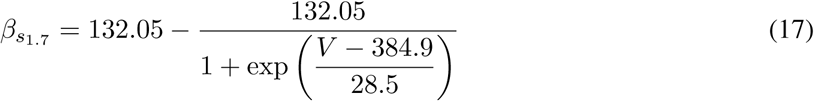

For *x* = *m*_1.7_, *h*_1.7_, *s*_1.7_,

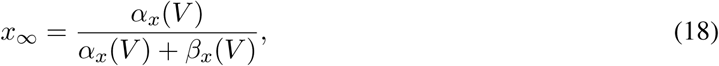

and

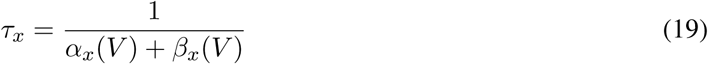

The equations for Na_v_1.7 were extracted from [29, 6]. *m*_1.7_ is the activation gating variable, *h*_1.7_ the fast-inactivation gating variable, and *s*_1.7_ the slow-inactivation gating variable.

#### Na_v_1.8 equations

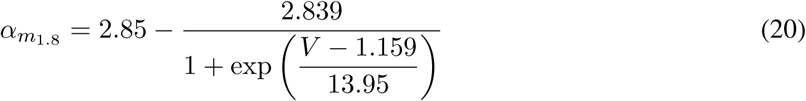

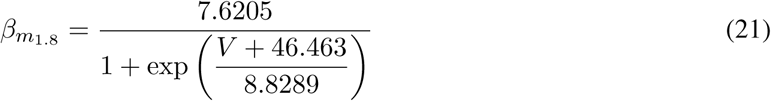

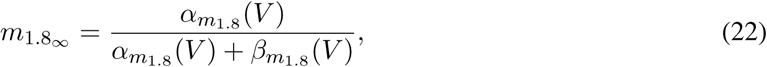

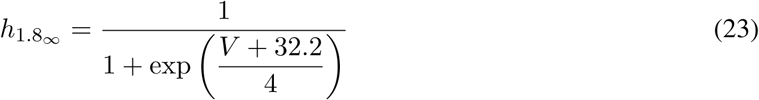

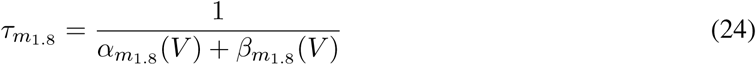

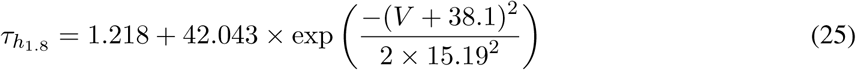

The equations for Na_v_1.8 were obtained from [29, 6]. *m*_1.8_ and *h*_1.8_ are activation and inactivation gating variables, respectively.

#### KDR equations

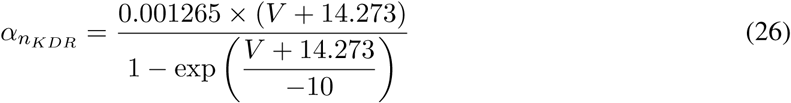

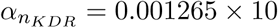 if *V* = −14.273.

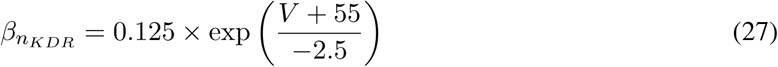

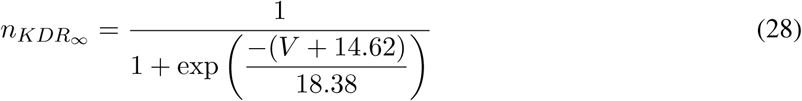

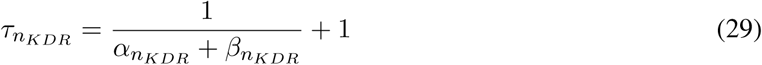

The equations of KDR channel were obtained from [27]. *n*_*KDR*_ is the activation gating variable. Inactivation gating variables are not present.

#### KA equations

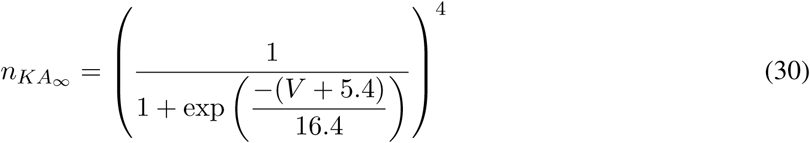

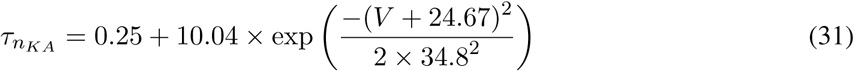

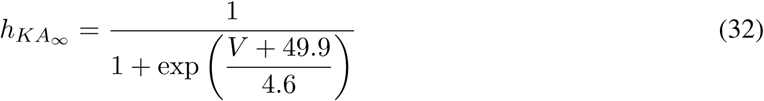

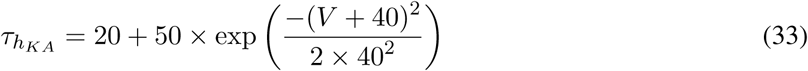

If 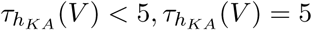.

The equations for KA channel were obtained from [29]. *n*_*KA*_ is the activation gating variable and *h*_*KA*_ is the inactivation gating variable.

#### Model parameter values

The model parameter values and the corresponding references are mentioned in Table 2.

### XPPAUT settings

All the bifurcation diagrams were primarily generated from XPPAUT [11]. Confirmation of the results were done using MATCONT [8]. Moreover, two parameter continuation was performed using MATCONT. All the plots were generated using MATLAB [21].

`NTST` = 100, `Method` = Stiff, `Tolerance` = 1e-07, `EPSL`, `EPSU`, `EPSS` = 1e-07, `ITMX`, `ITNW` = 20, `Dsmin` = 1e-05, `Dsmax` = 0.05 (there may be a need to adjust `Dsmax` to 0.1). There may be a need to adjust `Ds` to 0.01. All other settings were same as default.

### MATCONT settings

`MaxCorrIters` = 20, `MaxTestIters` = 20, `FunTolerance` = 1e-6, `VarTolerance` = 1e-6, `TestTolerance` = 1e-5, `MaxStepsize` = 0.01.

When keeping paclitaxel as the bifurcation parameter, following settings are changed from the above: `MaxStepsize` = 0.001, `InitStepsize` = 0.0001, `MinStepsize` = 1e-7.

### Reagents

200 nM A-803467 (Na_v_1.8 blocker) and 100 *µ*M L-alpha-phosphatidyl-D-myo-inositol 4,5-diphosphate, dioctanoyl (PIP_2_, KDR enhancer) were diluted in NbActiv4 recording media (BrainBits, Springfield, IL, USA). Complete saline solution (CSS) was made from 137 mM NaCl, 5.3 mM KCl, 1 mM MgCl_2_-6H_2_O, 25 mM sorbitol, 10 mM HEPES, and 3 mM CaCl_2_ equilibrated to pH 7.2.

### Primary Cell Culture

Dorsal root ganglia (DRG) was extracted from wild-type Sprague-Dawley rat pups 7-14 days old. Animals were maintained in the norovirus-negative facility of the Centrally Managed Animal Facilities at Purdue University. They were housed in at a constant temperature and humidity on a 12:12 light-dark cycle (lights on 0600-1800) with ad lib access to food and water according to as approved by the Purdue Animal Care and Use Committee and the Institutional Animal Care and Use Committee (IACUC) and will be conducted in accordance with the National Institutes of Health Guide for the Care and Use of Laboratory Animals. Pups were placed on a paper towel on ice for two minutes before decapitation. The spinal cord was extracted and the DRG was dissected and placed in complete saline solution (CSS). Cells were prepared by centrifuging at 2.5 g for 30 seconds then adding 1.5 mg*/*mL of collagenase A in CSS with 0.05 mM EDTA. After rotating in the 37C incubator for 20 minutes, cells were spun at 2.5× g for 30 seconds. 1.5 mg*/*mL collagenase D and 30U papain in CSS were added then placed in the incubator rotator for 20 minutes. Cells were spun at 2.5× g for 3 minutes. DRG were triturated in 1 mL of 0.15% trypsin inhibitor and 0.15% bovine serum albumin (BSA) in Dulbecco’s Modification of Eagle’s Medium (DMEM) medium with 10% fetal bovine serum (FBS) (Corning, Corning, NY, USA) then spun again at 2.5×g for three minutes before placing in a 40*µ*M filter before seeding on a coated plate. DRG from one pup was divided into six wells. Culture media was changed after two days to NbActiv4 recording media (BrainBits, Springfield, IL, USA). Four days after seeding, culture was recorded.

### Micro/multielectrode Array (MEA)

Firing properties was recorded using a Maestro Pro (Axion BioSystems, Atlanta, GA, USA). Twelve well plates of 64 electrodes per well used for culture. Plates were coated the day of seeding by incubating with poly-d-lysine for two hours, washing with sterile milli-q water three times, then incubating with laminin for one hour. All recordings were at 37°C. The plate was recorded before treatment and 24 hours after treatment. Then 200 *µ*L of 1.0 *µ*M capsaicin was added and recorded for two minutes as a positive control. Analysis was performed using the manufacturer’s software, Axion BioSystems Integrated Studio (AxIS) and NeuroMetric Tool. An electrode (n=1) was considered active if there was more than two action potentials in the baseline, response to buffer, or response to capsaicin. Mean firing rate (Hz) was calculated for active electrodes. An electrode will be considered active (n=1) if it has >1 spikes per 200 seconds. Chili pepper compound capsaicin will be added into the system to trigger a sensory response. Fold change of response to treatment was calculated based on baseline of the culture before treatment (treatment after 24 hours)/(cells at time zero before treatment). Recordings were compiled from different cultures extracted from different animals on different days.

### Statistics

A Shapiro-Wilk test determined that the data was not normal so Krushal-Wallis and Mann-Whitney U test were used to determine statistical significance of treatment differences. Results are presented as mean ± S.E.M. Effect size is Cohen’s D. P-value was set at p* ≤ 0.05 (*), p ≤ 0.01 (**), and p ≤0.001 (***). P-value is the probability that the means of two groups are either the same or different based on a threshold level of marginal significance (p=0.05). If p<0.05, we are 95% confident that the means are significantly different. Furthermore, we use effect size (reported here as standard error of the mean and Cohen’s D in the supplemental) to show the magnitude of significance and confidence. Both use mean, sample size, and variance to provide statistical certainty that our sampled dataset reflects the observed phenomenon population as a whole. GraphPad Prism 8.3.0 was used to determine statistical significance.

## Supporting information

**S1 Fig.**

**S2 Fig.**

## Acknowledgments

M.E. is supported by the National Science Foundation (NSF) Graduate Research Fellowship Program (GRFP) (DGE-1842166). We thank Dr. Charlie Zhang for technical support in performing the MEA experiments, Dr. Haroon Anwar, New Jersey Institute of Technology, USA, for support with model selection and building, and Max Planck Institute for Dynamics of Complex Technical Systems, Germany, for sponsoring trips to foster discussions. This project was supported, in part, with support from the Indiana Clinical and Translational Sciences Institute funded, in part by Award Number UL1TR002529 from the National Institutes of Health, National Center for Advancing Translational Sciences, Clinical and Translational Sciences Award. The content is solely the responsibility of the authors and does not necessarily represent the official views of the National Institutes of Health.

